# Strong hybrid male incompatibilities impede the spread of a selfish chromosome between populations of a fly

**DOI:** 10.1101/248237

**Authors:** Rudi L. Verspoor, Jack M. L. Smith, Natasha L. M. Mannion, Gregory D. D. Hurst, Tom A. R. Price

## Abstract

Meiotically driving sex chromosomes manipulate gametogenesis to increase their transmission at a cost to the rest of the genome. The intragenomic conflicts they produce have major impacts on the ecology and evolution of their host species. However their ecological dynamics remain poorly understood. Simple population genetic models predict meiotic drivers will rapidly reach fixation in a population and spread across a landscape. In contrast, natural populations commonly show spatial variation in the frequency of drivers, with drive present in clines or mosaics across species ranges. For example, *Drosophila subobscura* harbours a Sex Ratio distorting drive chromosome (“SR*s*”) at 15-25% frequency in North Africa, present at less than 2% frequency in adjacent Southern Spain and absent in other European populations. Here, we investigate the forces preventing the spread of the driver northward. We show that SR*s* has remained at a constant frequency in North Africa, and failed to spread in Spain. We find strong evidence in favour of our first hypothesis, genetic incompatibility between SRs and Spanish autosomal background. When we cross SR*s* from North Africa onto Spanish genetic backgrounds we observe strong SR*s* specific incompatibilities in hybrids. The incompatibilities increase in severity in F2 male hybrids, leading to almost complete infertility. We find no evidence supporting a second hypothesis, that there is resistance to drive in Spanish populations. We conclude that the source of the stepped frequency variation is genetic incompatibility between the SR*s* chromosome and the genetic backgrounds of the adjacent population, preventing SR*s* spreading northward. The low frequency of SR*s* in South Spain is consistent with recurrent gene flow across the Strait of Gibraltar combined with selection against the SR*s* element through genetic incompatibility. This demonstrates that incompatibilities between drive chromosomes and naïve populations can prevent the spread of drive between populations, at a continental scale.

## Introduction

Meiotically driving chromosomes are a class of selfish genetic element that spread through the selective destruction of gametes that bear the homologous chromosome (Burt and Trivers, 2006). This destruction ensures representation of the driving chromosome in greater than 50% of gametes of heterogametic parents, which in an XY system is the male. The drive will then result in particularly profound increases in the frequency of the element in species where females mate just once, as here within-ejaculate sperm competition is particularly strong. Drive phenomena are conceptually important as examples of intragenomic conflict, but are also studied as important ecological and evolutionary forces (Lindholm et al., 2016, Jaenike, 2001). They have been considered important in the evolution of sex ratios (Hamilton, 1967), mating rate (Price et al., 2008), in generating coevolutionary cycles with autosomal resistance genes (Bastide et al., 2011), and in the generation of reproductive isolation and hence speciation (Johnson, 2010, Phadnis and Orr, 2009).

The frequency of driving chromosomes is commonly heterogeneous over space (Lindholm et al., 2016). This variation takes a variety of forms. In *Drosophila pseudoobscura*, the driving SR X-chromosome shows clinal variation in the USA, being present at low frequency in Northern populations compared to Southern (Sturtevant and Dobzhansky, 1936). The frequency of driving X-chromosomes in *D. simulans* is a geographical mosaic, with high frequency in some populations, medium in others, and absence in some (Bastide et al., 2011). In the house mouse, populations also harbour the autosomal driving t-haplotype at varying frequencies (Lenington et al., 1988). Past work on *D. subobscura* indicates a stepped change in the frequency of the Sex Ratio distorting drive chromosome (henceforth referred to as “SR*s*”) between North Africa and Southern Europe, with SR*s* present at 15-25% frequency in North African samples, but at 0-2% in adjacent Southern European populations (Jungen, 1967, Hauschteckjungen, 1990, Prevosti, 1974).

Some of the causes of this spatial heterogeneity are known in some cases. For *D. pseudoobscura* and *D. neotestacea*, drive frequency is negatively associated with female remating rate, compatible with a reduction in intra-ejaculate competition (Price et al., 2014, Pinzone and Dyer, 2013). Female remating is also proposed to play a role in preventing the spread of the driving *t-haplotype* in house mice (Sutter and Lindholm, 2015, Manser et al., 2011). In *D. simulans*, variation in resistance gene presence is important (Bastide et al., 2011). Modification of multiple meiotic drive types by Y-linked suppression also exhibits geographic heterogeneity in *D. paramelanica* (Stalker, 1961). Restrictions of abiotic conditions have also been proposed to influence drive frequency in *D. neotestacea* (Dyer, 2012). In *D. subobscura*, however, the causes of differences in frequency across its range remain uncertain.

Here, we examine the causes of variation in the frequency of the SR*s* chromosome in *D. subobscura*. SR*s* was first recovered and characterized as a case of sex chromosome meiotic drive in *D. subobscura* collected from Tunisia in the sixties, where it had reached a frequency of 15-25% (Jungen, 1967, Jungen, 1968). Later studies in Southern Europe and Morocco between 1974 and 2002 revealed the SR*s* chromosome type was present in Morocco at 5-25% frequency, in Southern Spain at 0-2% frequency, and was absent in Italian, French and Northern Spanish populations (Prevosti, 1974, Sole et al., 2002). Variation in female mating rate can be excluded as a cause of SR*s* frequency heterogeneity, as the species is monandrous in the populations tested for SR*s* (Verspoor et al., 2016). Based on earlier work by Hauschteck-Jungen (1990), two alternate explanations for the spatial heterogeneity observed can be proposed. First, the SR*s* chromosome may be incompatible on genetic backgrounds outside North Africa. In support of this, a cross where an SR*s* chromosome from Tunisia was placed onto the genetic background of a Swiss isoline resulted in infertile males (Hauschteckjungen, 1990). If this incompatibility with SR*s* were to be similarly observed across diverse Southern Spanish genetic backgrounds, it could explain the stepped change in SR*s* frequency across the Straits of Gibraltar – SR*s* would be introduced by migration but then not spread. The second hypothesis to account for the failure of SR*s* to spread northwards is that there may be genetic resistance to the drive action in Southern European populations. Here, SR*s*/Y individuals are viable and fertile on a Southern European background, but drive does not occur and thus SR*s* does not spread.

In this paper, we first assess the frequency of SR*s* in North African and southern Spanish populations today, to determine whether the stepped change in frequency is still present, and thus represents an equilibrium (rather than transient) condition. We then test the two hypotheses for explaining the rarity of SR*s* in Southern Spain. First by examining if the SR*s* chromosome is compatible with a Spanish genetic background, and second by testing whether there is any evidence of resistance to drive in Southern Spain. Our results suggest that the failure of SR*s* to establish in Southern Spain is caused by the presence of incompatibilities between SR*s* and the Spanish genetic background, with SR*s*/Y individuals have greatly reduced fitness in the presence of Spanish autosomes (whilst non-driving X-chromosomes from North Africa are compatible). We hypothesize that it is intragenomic conflicts between SR*s* and autosomes that have driven this genomic incompatibility, which may therefore be a case of incipient conflictual speciation.

## Methods

### Wild fly collections and establishing current frequency of SR*s* phenotype

Flies were collected from three populations; Tabarka, Tunisia (36.57_N, 8.45_E) and Punta Umbria, Spain (37.10_N, 6.57_W) in April 2013 and Amizmiz, Morocco (31.19_N, 8.25_W) in April 2016 (Verspoor et al., 2015). Flies were caught from wild populations using banana, yeast and beer baits for collections at all three locations (Markow and O’Grady, 2005). Wild-caught females were brought into the laboratory and their offspring highly inbred to create isofemale lines (David et al., 2005), which captures wild genotypes and minimizes adaptation to the laboratory. Wild caught males were mated to a laboratory female to measure the sex-ratio of the offspring they produce. One Tunisian male was also used to isolate the SR*s* driving chromosome to be used in hybrid crosses (Verspoor et al., 2016).A sex-ratio of >85% females was used to assign status of SR*s* to a male (Hauschteckjungen, 1990).

### Investigating hybrid incompatibilities that could impede the spread of SR*s*

#### Testing compatibilities of SRs vs non-SRs chromosomes in native and naive populations

We searched for costs associated with the selfish SR*s* chromosome by testing the fitness of three types of X-chromosomes (a driving SR*s* X-chromosome, a selection of non-driving X-chromosomes from Tunisia and a selection of non-driving X-chromosomes from Spain) on different genetic backgrounds (either 100% from the population of origin (Native), or a 50:50 mix of native and foreign backgrounds). For details of collections of isolines see (Verspoor et al., 2015). SR*s* males from Tunisia, and non-driving males from both Tunisia and Spain were crossed to an outbred population from a selection of wild isolines (home vs away). This produced experimental males with either their full native or 50% non-native genetic background (Supplementary figure 1a). For each treatment, 80 replicates were set up. To test if the population of origin of the female was important, males were mated to either a female from Spain or from Tunisia, and this was tested as a separate factor. All flies used were seven days old, to ensure sexual maturity (Holman et al., 2008). Males were paired individually to females for seven days to lay eggs. The number of offspring was counted and analysed using ANOVA and Tukey’s post hoc tests. Offspring sex ratio was used to confirm male X-chromosome genotype (>85% female = SR*s*). Analyses were carried out in R (R Development Team, 2011).

**Figure 1.**
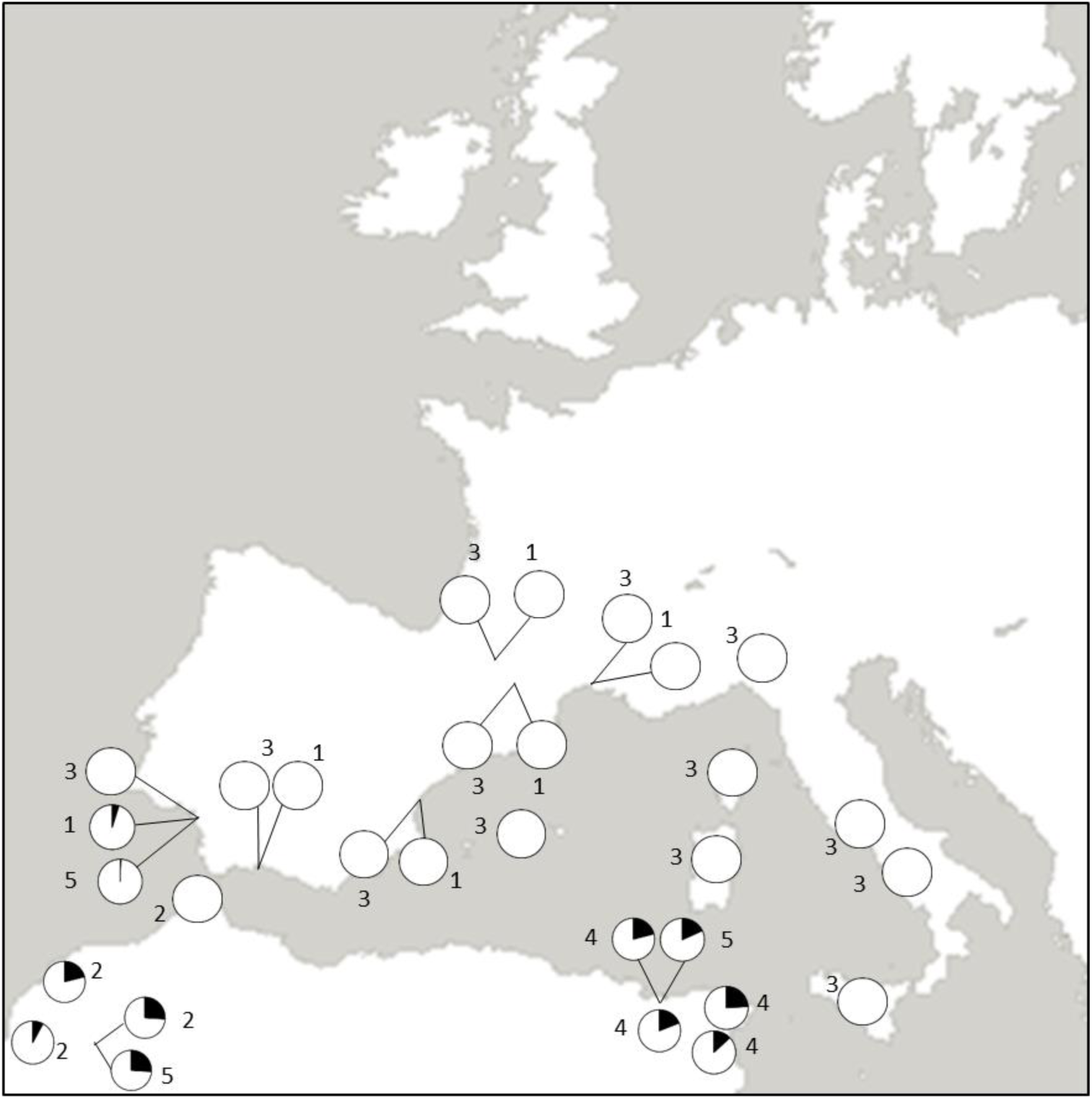
Map of driving and non-driving X-chromosomes across Southern Europe and North Africa. Pie charts show the proportion of SR*s* (in black) and non-driving (white) X-chromosomes. The numbers represent the years and sources of the collections (1 – 2002 (Sole et al., 2002), 2 – 1974 (Prevosti, 1974), 3 – 1984 (Prevosti et al., 1984), 4 – 1968 (Jungen, 1968), 5 – 2013-15 collections Supplementary table 1).

#### Hybrid incompatibility of SRs males across three populations

The strength of the hybrid incompatibility of SR*s* was further tested by comparing its fitness across different isofemale lines from three populations (Tunisia, Spain and the UK). Each isofemale line (or "isoline") comprises of the highly inbred descendants of a single wild caught. Experimental males were produced by crossing SR*s* homozygote females to males from an isoline (Tunisia – 15 isolines; Spain – 16 isolines; UK; 8 isolines; supplementary figure 1b). These produced experimental SR*s* hybrid males with a 50% Tunisian and 50% Tunisia, UK or Spanish genetic backgrounds. From each of the 39 isolines, 40 SR*s* males were paired with a virgin female from an outcrossed Tunisian population. Pairs were mated and the number of offspring produced was recorded as described in the previous section. To avoid pseudo replication a mean number of offspring and sex-ratio was calculated for each isoline cross. From these, ANOVAs were used to test if there was a significant effect of population of origin on the number of offspring produced and the strength of drive. Analyses were carried out in R (R Development Team, 2011).

#### Fitness of the SRs chromosome in an F2 hybrid genetic background

We then tested the fitness of SR*s* when it had been introgressed for 2 generations into a foreign Spanish background. To produce the experimental males we crossed F1 heterozygote females carrying one SR*s* X-chromosome to males from three randomly selected Spanish isolines. The resulting male offspring now carried either an SR*s* or a Spanish non-driving X-chromosome with a ~25% Tunisia/75% Spain genetic background. These focal males were mated as described above to a Spanish female, and the number of offspring produced was recorded. Focal males that produced fewer than 5 offspring could not be reliably assigned by offspring sex ratio, and so their X-chromosome type was confirmed by sequencing the *G6P* gene region in both directions and using SNPs to identify the chromosome of origin (see SOM for details). The number of offspring produced was analysed using a Wilcoxon rank test as it was not possible to transform the data to meet the requirements of parametric tests.

#### Testing for rescue of SRs phenotype by backcrossing to the Tunisian genetic background

The F2 SR*s* hybrid males were almost entirely infertile, producing only a few female F3 offspring. To examine whether SR*s* fertility could be rescued by increasing the proportion of the background that was Tunisian, we crossed half of these females to a random male from an outbred Tunisian population, and half to an outbred Spanish population. As the focal females were heterozygotes, carrying one SR*s* chromosome, half of their sons would be expected to carry SR*s*. Male offspring of each focal female were mated as above. Males were subsequently assigned to three phenotype categories: SR*s* if the sex ratio of their offspring was >85% female, non-driving if the sex ratio was 50:50, and unknown if they produced 5 or fewer offspring. If Tunisian autosomes do rescue SR*s*, then the SR*s* phenotype would only appear in the backcross to Tunisian males, as SRs would remain sterile in the Spanish background.

#### Fitness of Tunisian SRs background in Moroccan genetic backgrounds

As a separate follow up experiment we tested if there were incompatibilities in a different population from within North Africa. The Tunisian SR*s*, was crossed into 11 different isolines from Amizmiz, Morocco, using the same crossing approach as when it was crossed into Spanish genetic backgrounds to generate experimental males. These males are 50/50 Tunisia/Morocco hybrids, and carry a range of genetic backgrounds from the natural population in Morocco (Supplementary figure 7.1). SR*s* on a Tunisian background was used as a control for comparison of the strength of drive and offspring production. For each isoline and the Tunisian control 25 replicates were set up to measure the number of offspring produced. Each of the Tunisian/Morocco SR*s* bearing males, and the pure Tunisian SR*s* males, were mated to a virgin Tunisian female collected from an outbred mass population originating from a mix of Tunisian isolines. Lines were then compared against each other using an ANOVA.

### Testing for evidence for suppression of the SR*s* drive phenotype

#### Sex-ratio distortion of SRs males across multiple populations

If the rest of the genome has evolved resistance against drive, we expect genetic variation in the sex ratios produced by males carrying SR*s*. We tested whether the strength of the SR*s* sex-ratio distorting phenotype differed when it was in hybrid genetic backgrounds between different populations. For details of the F1 Experimental males see supplementary figure 1b. Pairs were mated as described in the first methods section and the offspring produced over 7 days were sexed and counted. Proportion of female offspring was transformed using an arcsine transformation before analysis. A mean sex ratio for each line was calculated and populations were compared using ANOVA and Tukey’s post hoc tests.

#### Fertility and Y-chromosome status of sons of SRs males

In some systems, sons of meiotic drive males are actually infertile pseudomales, and so do not represent effective suppression of drive (Cobbs, 1992). To check the few males produced from SR*s* fathers were not pseudomales, a subset of males offspring produced by males with the SRs chromosome and a Tunisian genetic background were tested for fertility by pairing them to two random virgin females from an outcrossed Tunisian population. After mating the pseudomales as described in section 1 of the methods, vials were checked for larval action to confirm the males were fertile. These same males were then assayed for the presence of a Y-chromosome using the *kl2* marker (Herrig et al., 2014). The presence of a Y-chromosome was confirmed using gel electrophoresis, with a positive and negative control.

## Results

### SRs frequency remains consistent in North Africa and Spain

We found evidence for moderate levels of the SR*s* phenotype in males screened from Morocco (18%) and Tunisia (12%) (Figure 1, Supplementary table 1). We found evidence for the SR*s* phenotype at very low frequencies in males in South Spain (0.5%) (Figure 1, Supplementary table 1).

### Strong incompatibilities reduce SR*s* fitness on naïve genetic backgrounds

#### The compatibility of SRs vs non-SRs chromosomes in native and naive populations

Female origin had no impact on offspring production or sex ratio (ANOVA F1,418=0.474 p=0.492), so this factor was removed first from the ANOVA. We established a source of selection against the SR*s* element by examining the fitness of males carrying SR*s*, non-driving North-African, or non-driving Spanish X-chromosomes on their native and hybrid backgrounds (Figure 2A). We found a highly significant interaction between X-chromosome type and genetic background (Figure 2A; ANOVA: F2,419 = 30.64 p <0.001). Males that carried a non-driving X-chromosome showed equally high levels of fitness, irrespective of whether they were on a Spanish, Tunisian, or mixed genetic background (Tukey’s post hoc test: P>0.861 in all comparisons, Supplementary table 2). The fitness of SR*s* males that had a Tunisian genetic background equalled the high fitness of non-driving males (Tukey’s post hoc test: P>0.875, Supplementary table 2). However, males that carried SR*s* on a mixed Spanish/Tunisian genetic background produced fewer than half the number of offspring than all other male types (Tukey’s post hoc test: P<0.001 in all comparisons, Supplementary table 2).

**Figure 2.**
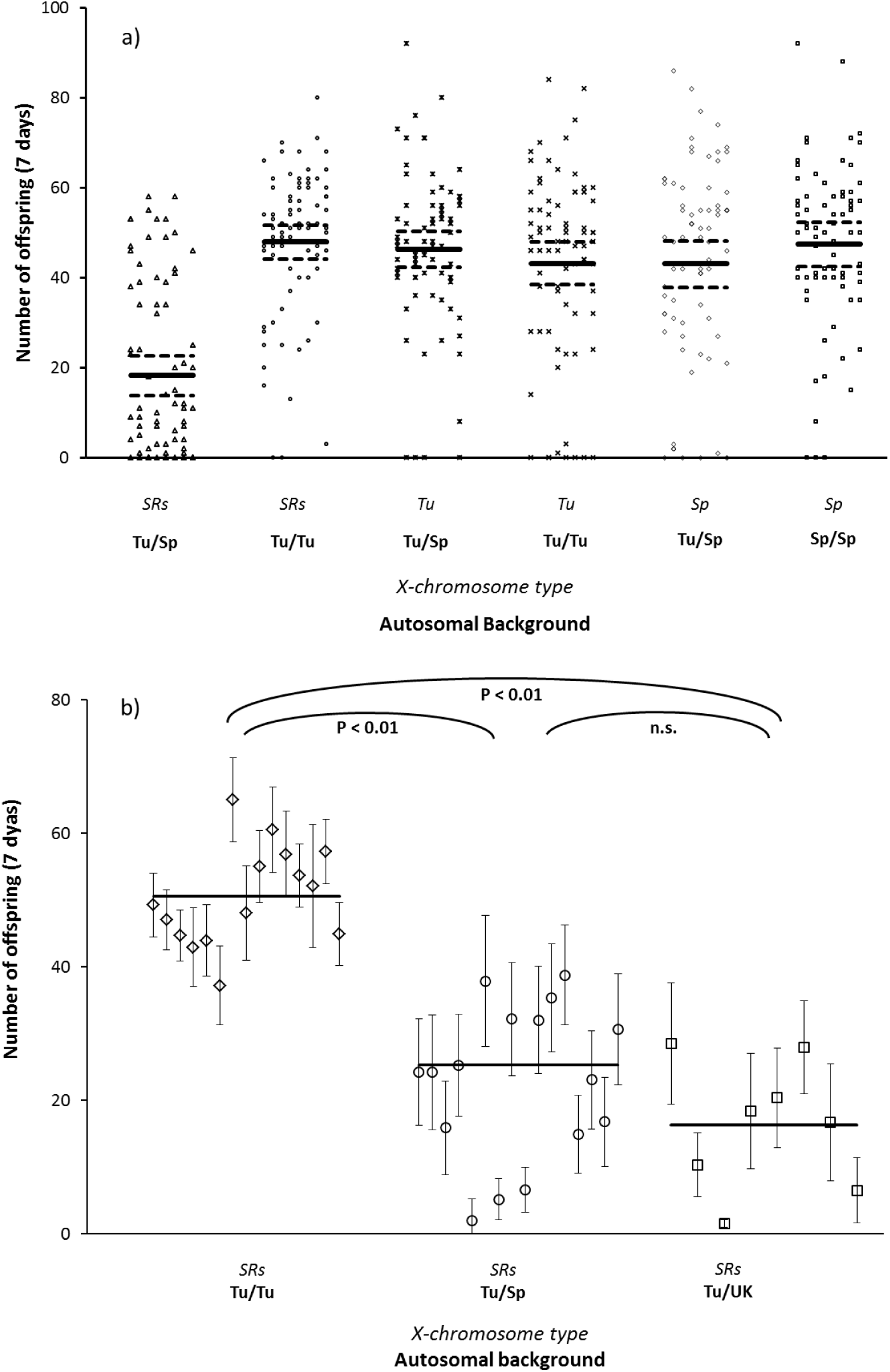
a) The number of offspring produced by three different categories of X-chromosome (*SRs*, non-driving Tunisian – *Tu*, non-driving Spanish – *Sp*) on different autosomal backgrounds (100% native autosomes or 50% foreign autosomes). Reduced fitness only occurs when SR*s* chromosomes occur in a hybrid background. Solid and dashed lined show the means and two SEM respectively. b) Mean number of offspring produced by SR*s* males with different genetic backgrounds, showing low fitness on hybrid backgrounds. Each point indicates the mean and 95% confidence intervals for a single isoline. Main lines show population means.

#### Hybrid incompatibility of SRs males across three populations

We crossed SR*s* into a number of isofemale lines from the native Tunisian population, the neighbouring Spanish population, and a distant UK population. There was equally strong evidence of incompatibility when the SR*s* chromosome is expressed on F1 hybrid backgrounds from both neighbouring (Spain) and distant (UK) populations (Figure 2B; ANOVA: F2,36 = 43.91 p < 0.001; see Supplementary Table 3). This result corroborates the previous evidence of strong incompatibilities between in Spanish backgrounds and SR*s*, and extended these incompatibilities to a more distant population sample from the UK.

#### The fitness of the SRs chromosome in an F2 hybrid genetic background

Expressing the SR*s* chromosome on an increasingly Spanish genetic background in an F2 backcross resulted in almost complete infertility of SR*s* males. Over ninety percent of these males produced fewer than five offspring, compared to Spanish X-chromosomes which show normal offspring production (Wilcoxon rank: n=176 W = 6958 p < 0.001; Figure 3). This result taken in concert with the previous two results show that the incompatibility caused by the SR*s* chromosomes increases when expressed on an increasingly Spanish genetic background from the F1 generation to the F2 generation.

**Figure 3.**
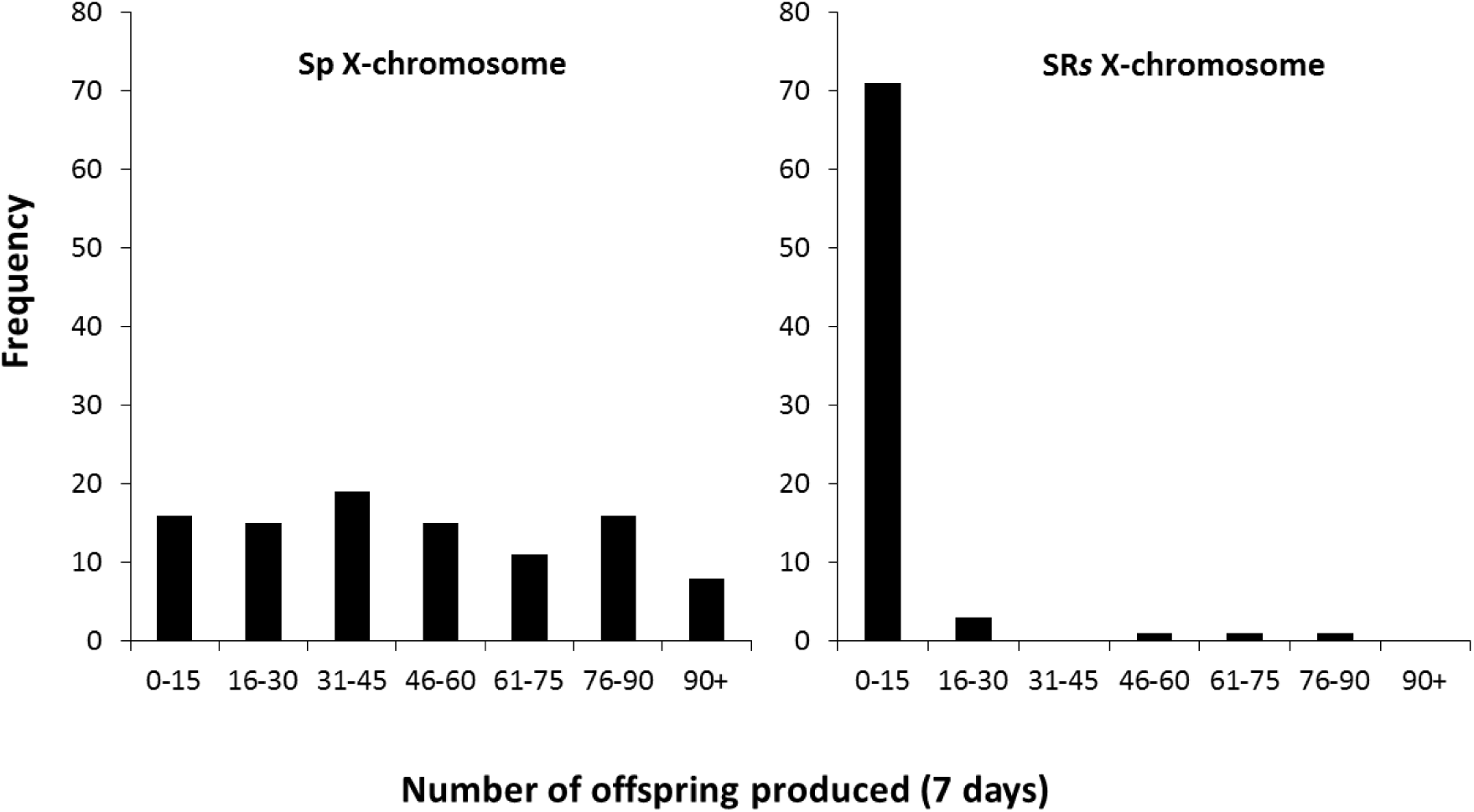
Histogram of offspring production for males, which were introgressed into the Spanish population for two generations, carrying the Spanish (Sp) X chromosome or SR*s*. The identity of the X-chromosome the males were carrying was identified using the SNP variation in the G6P locus (supplementary information 1).

#### Testing for rescue of SRs phenotype by backcrossing to the Tunisian genetic background

We tested if reintroducing Tunisian genetic material could rescue the fertility of the SR*s* chromosome by backcrossing a small number of F3 offspring that were produced to Tunisian males. The resulting F4 SR*s* sons had restored fertility (Supplementary Table 4). This restored fertility was specific to those crosses that reintroduced Tunisian genetic backgrounds.

#### The fitness of Tunisian SRs background in Moroccan genetic backgrounds

The result of the experiment crossing Tunisian SR*s* into Moroccan genetic backgrounds demonstrates that the hybrid incompatibility does not occur when the Tunisian SR*s* is crossed into genetic backgrounds from a population in Morocco. Hybrid crosses between Moroccan isolines and the Tunisian driver did not differ significantly in the number of offspring they produced when compared to a fully Tunisian background (ANOVA F11,245, p = 0.318; figure 4). There were no significant differences between any of the isolines using Tukey’s post-hoc tests (all p > 0.3).

**Figure 4.**
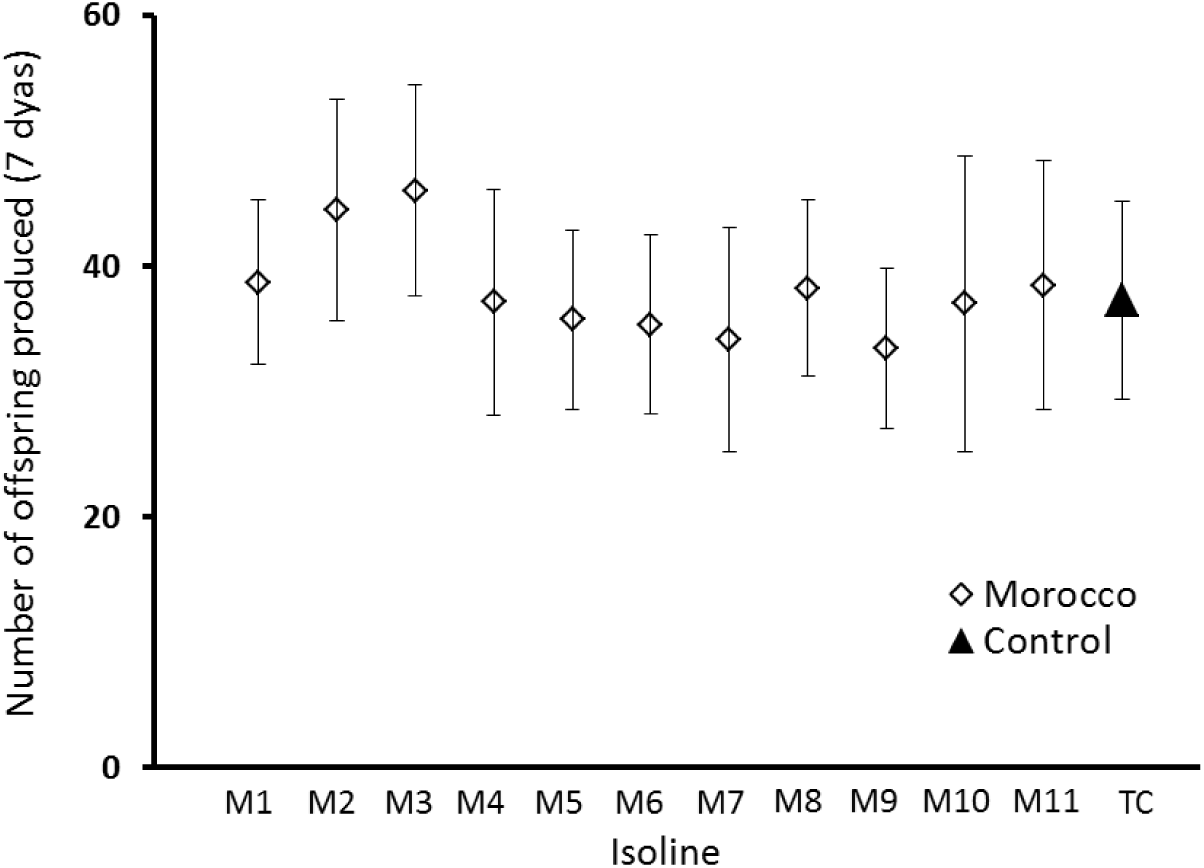
Total offspring produced by hybrid Moroccan/Tunisian males carrying the Tunisian SR*s* (diamonds) and pure Tunisian control males carrying SR*s* (triangle). Points indicate the mean number of offspring produced, while error bars indicate 95% confidence intervals.

### No evidence for suppression preventing the spread of drive

#### Sex-ratio distortion of SRs males across multiple populations

We compared the strength of drive in three different populations, Tunisia, Spain and the UK. We observed that drive is stronger in hybrids with partial Spanish and UK genetic backgrounds than in the Tunisian background (ANOVA F2,36=17.71 p < 0.001; figure 5). This is being driven by Tunisia showing evidence of suppression (Tukey’s post hoc test: P<0.0.01 in both comparisons, Supplementary table 5). Figure 5 also highlights that there are differences in the strength of drive between genetic backgrounds from Tunisia that are consistent between isolines. In contrast, the strength of drive appears to be consistently strong in all genetic backgrounds from Spain and the UK.

**Figure 5.**
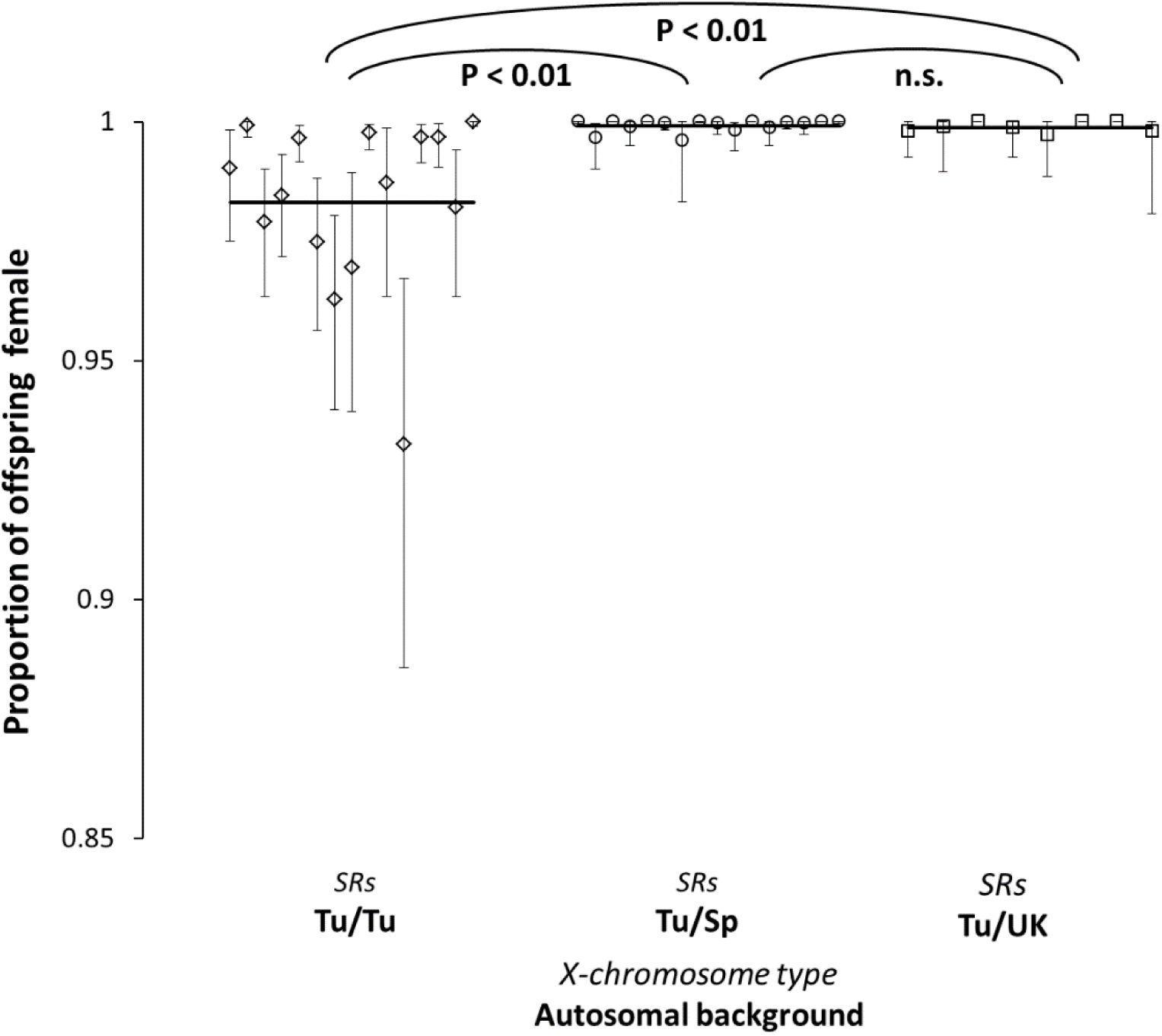
Offspring sex ratio of SR*s* males on native and Spanish/Tunisian and UK/Tunisian hybrid backgrounds. Each point indicates the mean and 95% confidence intervals for a single isoline. Main lines show population means.

#### Fertility and Y-chromosome status of sons of SRs males

When we examined a random selection of the male offspring from lines showing weak suppression, we found that all but one of these males was found to be both fertile and to carry a Y-chromosome. This result demonstrates that in this system there is true suppression in North Africa, as all but one of the sons produced by SR*s* fathers were fully fertile males carrying a Y chromosome (Supplementary figure 2) with only one pseudomale (Cobbs, 1992). This demonstrates that the Y-chromosome was succeeding in being transmitted to these few male offspring and that these males do produce functional sperm.

## Discussion

Meiotically driving chromosomes can have important evolutionary and ecological consequences (Lindholm et al., 2016). In a number of these systems, the driving chromosomes occur at different frequencies across species ranges (Lindholm et al., 2016, Jaenike, 2001). In some cases, factors underlying these differences have been identified, for example remating rates (Price et al., 2014, Pinzone and Dyer, 2013) or genetic suppression (Bastide et al., 2011, Stalker, 1961). However, these explanations are by no means ubiquitous. Thus, our broader understanding of factors that dictate the frequency of selfish driving chromosomes in natural populations remains incomplete.

In this study we aimed to further understand the causes of the difference of frequency in SR*s*, a driving X-chromosome, in the monandrous fruit fly *Drosophila subobscura*. Our field collections of *D. subobscura* from three populations (Tunisia, Morocco, and Southern Spain) confirm that the SR*s* phenotype is still present in all three locations (Figure 1). Frequencies of SR*s* were similar to previous samplings from Tunisia and Morooco (Jungen, 1967, Hauschteckjungen, 1990, Prevosti, 1974). However, in southern Spain we found the drive phenotype at slightly lower frequencies than previous reports (Sole et al., 2002). Overall, these results are consistent with the polymorphism of SR*s* across the range being roughly stable over the last 50 years.

What prevents SR*s* from increasing in frequency in Spanish populations? Our principle finding is that the lack of introgression of SR*s* into Southern Spain is associated with severe genetic incompatibilities between SR*s* and Spanish genetic backgrounds. Motivated by previous findings of hybrid failure in a single SR*s* and a Swiss isogenic lineage (Hauschteckjungen, 1990), we tested whether hybrid failure commonly occurred for SR*s* on the genetic background of the adjacent population. We observed strong SR*s* hybrid incompatibilities when SR*s* is found in Spanish genetic backgrounds. This hybrid incompatibility is not found for non-driving X-chromosomes from Tunisia. This drive specific incompatibility thus represents a powerful impediment to the spread of SR*s* in Europe. In contrast, we find no evidence for genetic suppression in Spanish populations preventing the spread of SR*s* into Spain. These results demonstrate inter-population incompatibilities are a novel mechanism that can prevent the spread of a driving chromosome. In this case, strong hybrid incompatibilities specific to a driving chromosome are blocking it from spreading into south Spain.

What is causing the evolution of these incompatibilities? Population specific co-evolution driven by genetic conflict between drivers and suppressors is a plausible explanation for these incompatibilities (Johnson, 2010, Crespi and Nosil, 2013). The evolution of genetic suppression of selfish driving chromosomes has been observed in a number of other systems (for example Bastide et al., 2011, Stalker, 1961). In the SR*s* system, the weak suppression in North Africa supports this hypothesis, as evidence that genetic conflicts over suppression are ongoing. Our results are therefore consistent with the involvement of drive and suppression in causing these incompatibilities. However, the observations to date represent interactions between an entire SR*s* chromosome and Spanish autosomes: they do not causally link drive itself to the incompatibility. Future work will need to examine the mechanism underlying incompatibility in depth, and in particular the role of the driver (as opposed to linked variants) in causing incompatibility. If the driver is shown to be causally associated with incompatibility, it will represent strong evidence in support of the hypothesis that drive/autosome coevolution may drive the primary stages of reproductive isolation, as hypothesized by (McDermott and Noor, 2012).

Will this incompatibility barrier remain intact in the long term? Measuring the degree of gene-flow between North Africa and Europe will be important in order to determine this. Historically, *D. subobscura* was likely divided into sub-populations by glaciation. However currently there is admixture and gene-flow between these populations (Krimbas, 1993). In addition, this admixture is occurring on the back drop of a changing climate, where the genetic assemblages of southern populations of *D. subobscura* in Europe are moving northwards (Balanya et al., 2006). It is not known if this shift northward in population genetics is also occurring between North African and southern European populations. The extent to which North African genetic backgrounds are arriving and establishing into southern Spain will strongly affect the strength and severity of any incompatibilities that exist as a barrier to SR*s* establishing there.

Could incipient hybrid incompatibilities be present in other drive systems? Both between sub-species and sister species there is already strong evidence for drive loci being associated with incompatibilities (Phadnis and Orr, 2009, McDermott and Noor, 2012). We see no reason to expect drive specific incompatibilities to be restricted to these systems. Incompatibilities created by co-evolution between suppressors and drivers could plausibly exist in other systems. In many systems suppressors of drivers occur across populations (see Jaenike, 2001 for review). Equally, if these incompatibilities are caused by linked variants that are locked up in large driving inversions, large inversions are not unique to the SR*s* systems in *D. subobscura*. Driving chromosomes often have large inversions, which reduce recombination and creates linkage across large areas regions (for example Dyer et al., 2007, Babcock and Anderson, 1996).

The system allows us a unique opportunity to gain insights into the early origins of incompatibility associated with meiotic drive. It is likely the SR*s* system experienced a degree of historic subdivision between populations. Is some subdivision and barrier to gene-flow necessary? If so, other systems where species have restricted gene-flow due to climate history or geographic isolation may be candidates for the same process. However, in the case of SR*s* it is still unknown if this incompatibility evolved in the form of the accumulation of minor incompatibilities or as one or two single large contributing loci. Determining the number and age of the loci that are contributing to this incompatibility would prove informative to understanding how these incompatibilities initially begin to form. Excitingly, our observation is of incipient incompatibilities in process. For this reason, the system provides a window to understanding the formation of early incompatibilities between populations, and could help answer some of these fundamental questions of process.

## Acknowledgements

The authors would like to thank M Booth and P. Giraldo for help during the experiments and A Agren for helpful comments on the manuscript. This project was funded by a NERC grant (ref. NE/P002692/1).

## Author contributions

RV, TP and GH conceived and designed the study. RV, NM and JS carried out the experiments. RV and TP analysed the data. RV, TP and GH drafted the manuscript with later contribution from all authors

## Data Accessibility

All data will be deposited at datadryad.

---

**Supplementary figure 1.**
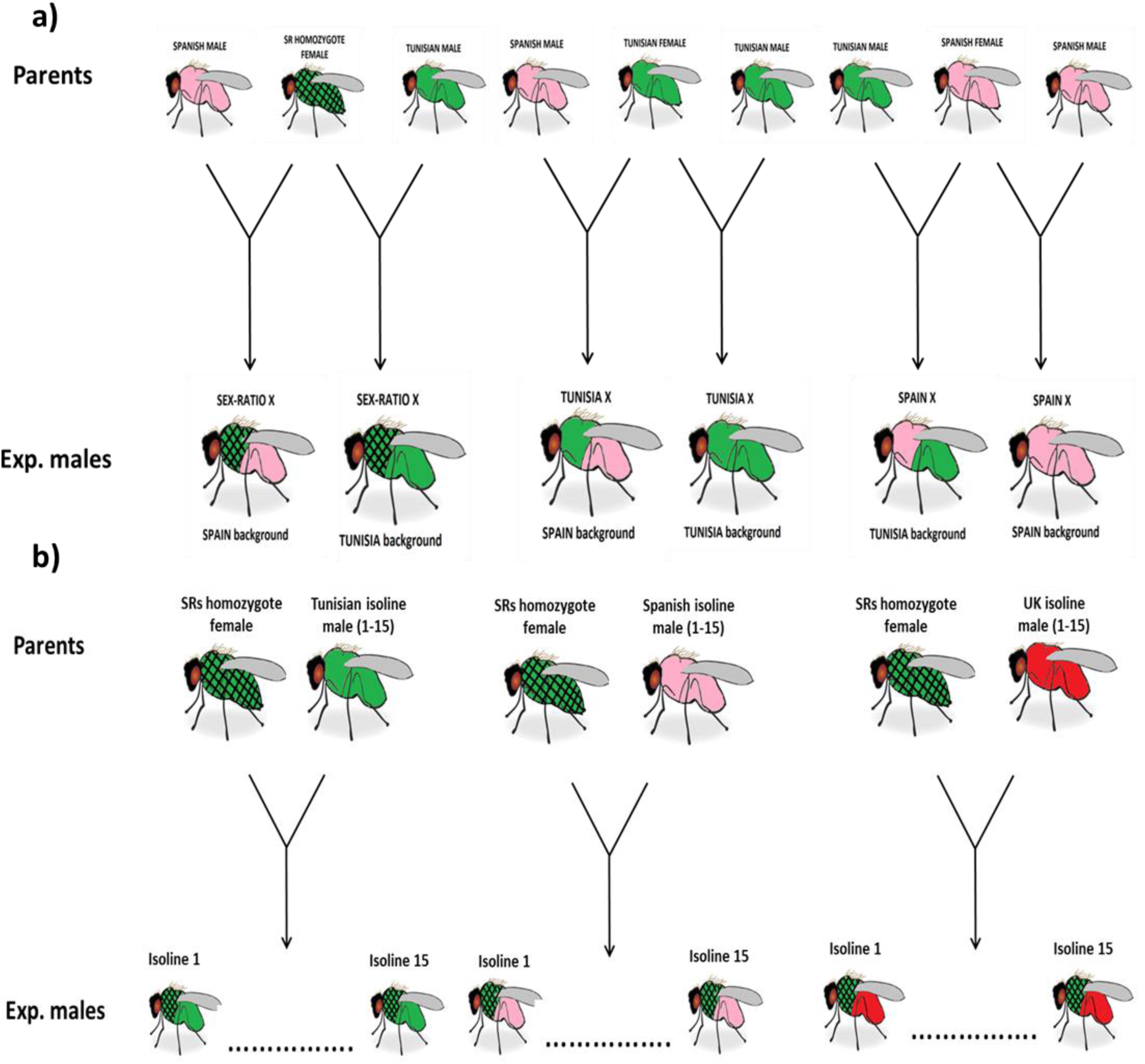
Showing the layout of the crossing schematics for a) Experiment 1 comparing the fitness of the SRs X-chromosome and non-driving X-chromosomes from Tunisia and Spain on native and hybrid populations genetic backgrounds b) Experiment 2 and 4 comparing the fitness costs of SRs and the levels of suppression of SRs in multiple isofemale lines across three populations. Colours indicate different genetic backgrounds from different populations. Checked pattern indicates the SR*s* X-chromosome is present in the cross.

**Supplementary figure 2.**
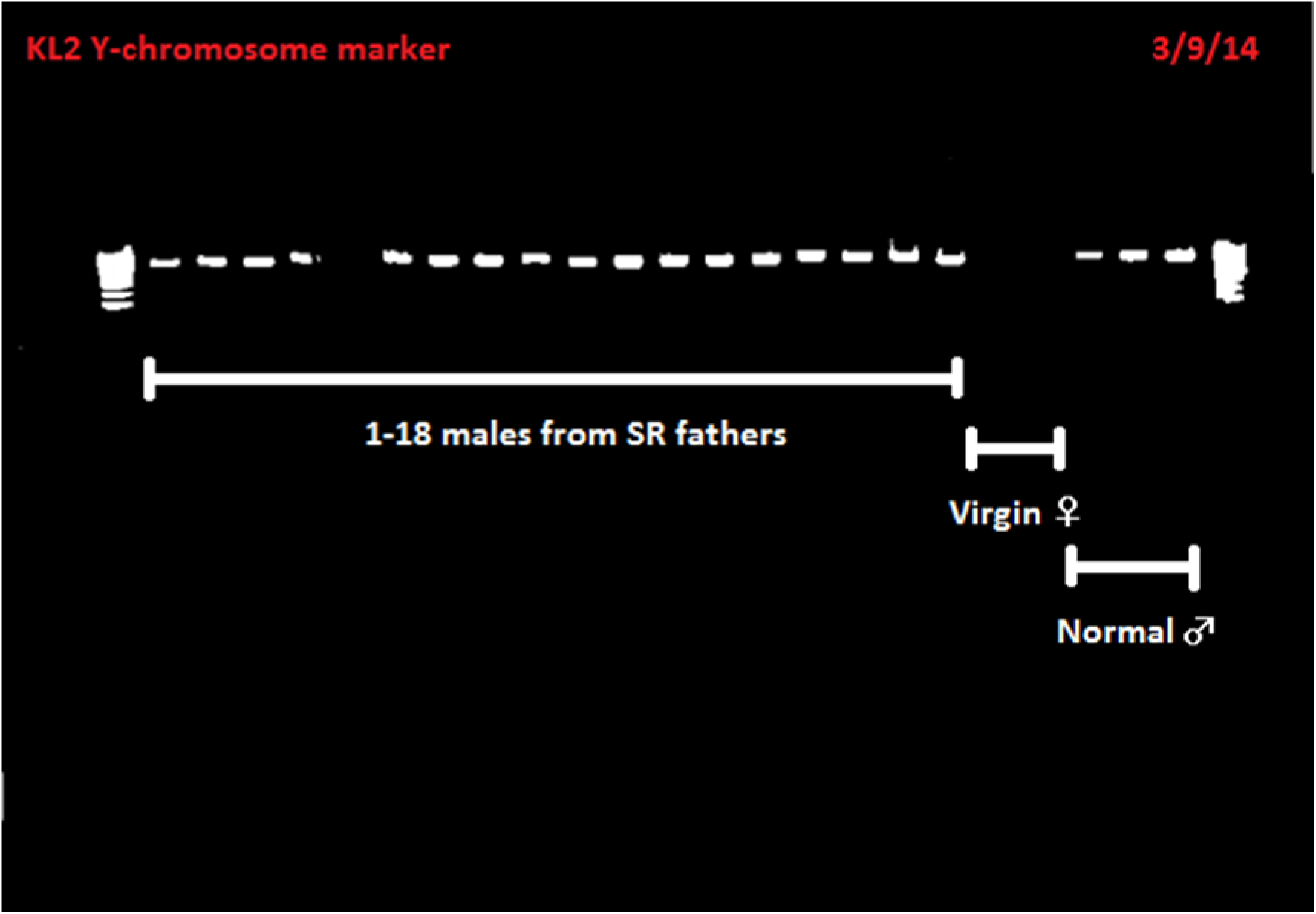
Gel electrophoresis image showing the amplification of the kl2 gene from the *Drosophila subobscura* Y-chromosome. PCR conditions were an initial 3min denaturing step, followed by 35 cycles of 94 for 30secs, 60 for 30 secs, 72 for 30secs, with a final elongation period of 10mins at 72. PCR products were determined using gel electrophoresis on a 1.5% agarose gel with 3μL Midori green per 100mL of TAE buffer. This image confirms that the few male offspring produced from SR*s* carrying males are carrying a Y-chromosome. All the males which carried a Y-chromosome were also found to be able to produce offspring when mated to a virgin female.

## Supplementary information 1 - G6P locus

The G6P locus, located on the X-chromosome (A chromosome) was used to differentiate SRs, from Spanish X-chromosomes. The forward and reverse primers used can be seen below (Forward primer – ATCATACCGCTCTGGATCTCAT, Reverse primer – GTGGAGCTGAGGATCTTGTTG). The reaction profile was an initial 3min denaturing step at 95°C, followed by 35 cycles of 95°C for 30secs, 60°C for 30 secs, 72°C for 30secs, with a final elongation period of 10mins at 72°C. PCR products were determined using gel electrophoresis on a 2.5% agarose gel with 3μL Midori green per 100mL of TAE buffer. For one of the Spanish X-chromosomes, there was no amplification so SRs was scored based on the presence of a PCR product. For the two remaining X-chromosome types Sanger sequencing was used to identify the X-chromosome by SNP variation. PCR products were cleaned using antarctic phosphatase and exonuclease 1, with an incubation of 45mins at 37°C followed by 15mintes at 80°C. Sequencing products were amplified using BigDye3.1 protocol with a sequencing program of 35 cycles of 96°C for 10secs, 50°C for 5secs, 60°C for 4mins. Sequencing was precipitated using 3M sodium acetate and cleaned with 70% ethanol. 10ul of Hi-Di formamide was then added and sequencing was carried out on and ABI3500xL genetic analyser. SNPs in the region were called using the software Geneious version 7.1.3.

Supplementary table 1 shows the proportion of female offspring produced from wild males caught in Tunisia and Spain collected in 2013. Males were each crossed to a 7 day old virgin female from their same population of origin. Males were conservatively classed as the SR*s* phenotype if they produced >85% female broods ((Hauschteckjungen, 1990).

| Prop. of offspring<br>female | Tunisia (n= 146) | Spain (n = 320) | Morocco (n=135) |
| --- | --- | --- | --- |
| 0.3-0.4 | 1 | 9 | 8 |
| 0.4-0.5 | 32 | 86 | 22 |
| 0.5-0.6 | 83 | 175 | 50 |
| 0.6-0.7 | 22 | 41 | 21 |
| 0.7-0.8 | 3 | 7 | 8 |
| 0.8-0.9 | 3 | 0 | 3 |
| 0.9-1.0 | 12 | 2 | 23 |

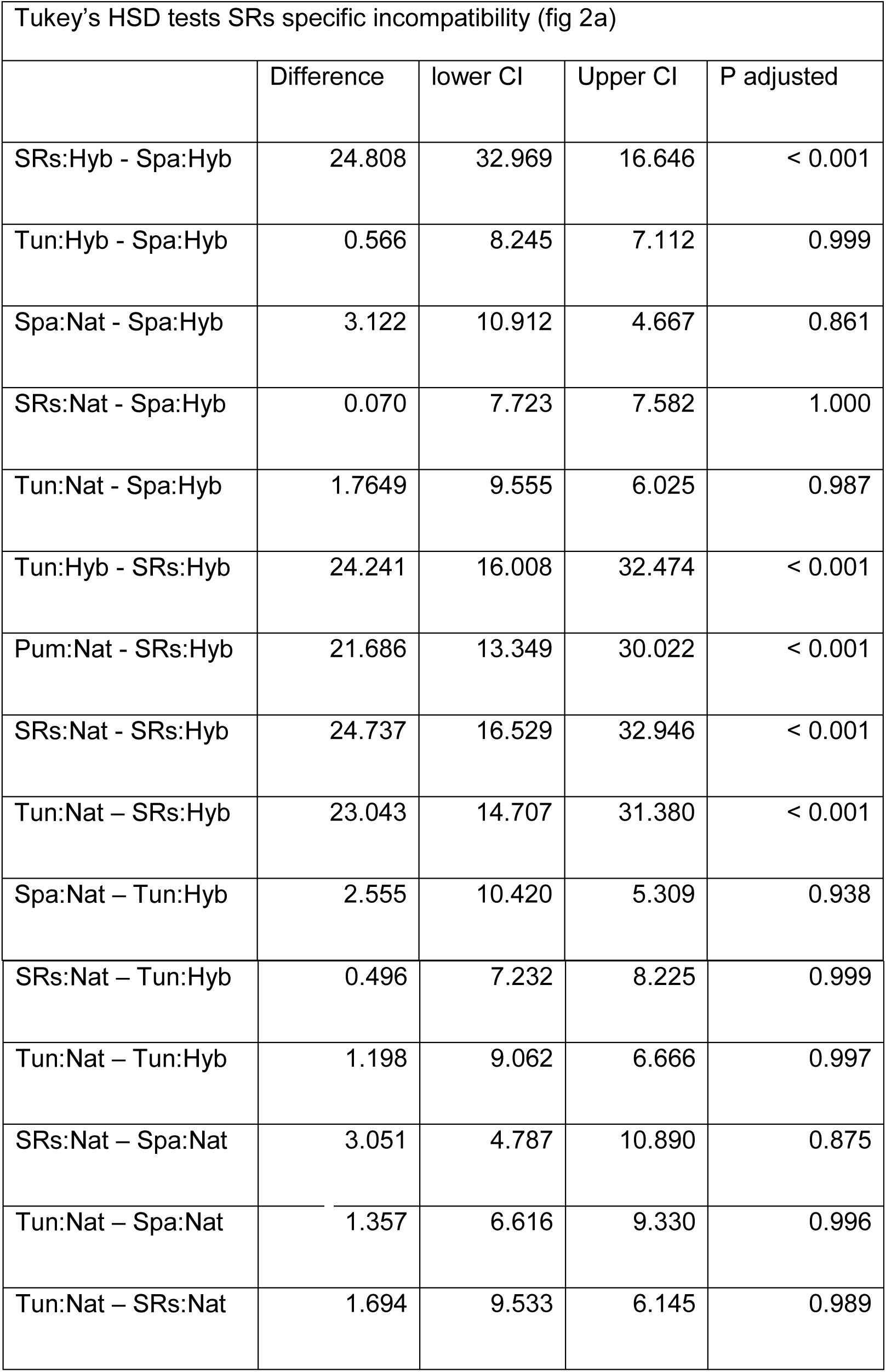
Supplementary table 2 showing Tukey’s post hoc tests on the differences in offspring produced by three types of X-chromosome (Driving SR*s* - "SR*s*", non-driving Tunisian – "Tun" and non-driving Spanish – "Spa") on two different population genetic backgrounds (100% their own native background – Nat or 50%/50% their own background and that of a different population – Hyb). The replicates for each category are as follows (SR*s*:Hyb n=61, SR*s*:Nat n=75, Spa:Hyb n=77, Spa:Nat n=70, Tun:Hyb n=73, Tun:Nat n=71)

| Tukey's HSD tests SRs specific incompatibility (fig 2a) |  |  |  |  |
| --- | --- | --- | --- | --- |
|  | Difference | lower CI | Upper CI | P adjusted |
| SRs:Hyb - Spa:Hyb | 24.808 | 32.969 | 16.646 | < 0.001 |
| Tun:Hyb - Spa:Hyb | 0.566 | 8.245 | 7.112 | 0.999 |
| Spa:Nat - Spa:Hyb | 3.122 | 10.912 | 4.667 | 0.861 |
| SRs:Nat - Spa:Hyb | 0.070 | 7.723 | 7.582 | 1.000 |
| Tun:Nat - Spa:Hyb | 1.7649 | 9.555 | 6.025 | 0.987 |
| Tun:Hyb - SRs:Hyb | 24.241 | 16.008 | 32.474 | < 0.001 |
| Pum:Nat - SRs:Hyb | 21.686 | 13.349 | 30.022 | < 0.001 |
| SRs:Nat - SRs:Hyb | 24.737 | 16.529 | 32.946 | < 0.001 |
| Tun:Nat – SRs:Hyb | 23.043 | 14.707 | 31.380 | < 0.001 |
| Spa:Nat – Tun:Hyb | 2.555 | 10.420 | 5.309 | 0.938 |
| SRs:Nat – Tun:Hyb | 0.496 | 7.232 | 8.225 | 0.999 |
| Tun:Nat – Tun:Hyb | 1.198 | 9.062 | 6.666 | 0.997 |
| SRs:Nat – Spa:Nat | 3.051 | 4.787 | 10.890 | 0.875 |
| Tun:Nat – Spa:Nat | 1.357 | 6.616 | 9.330 | 0.996 |
| Tun:Nat – SRs:Nat | 1.694 | 9.533 | 6.145 | 0.989 |

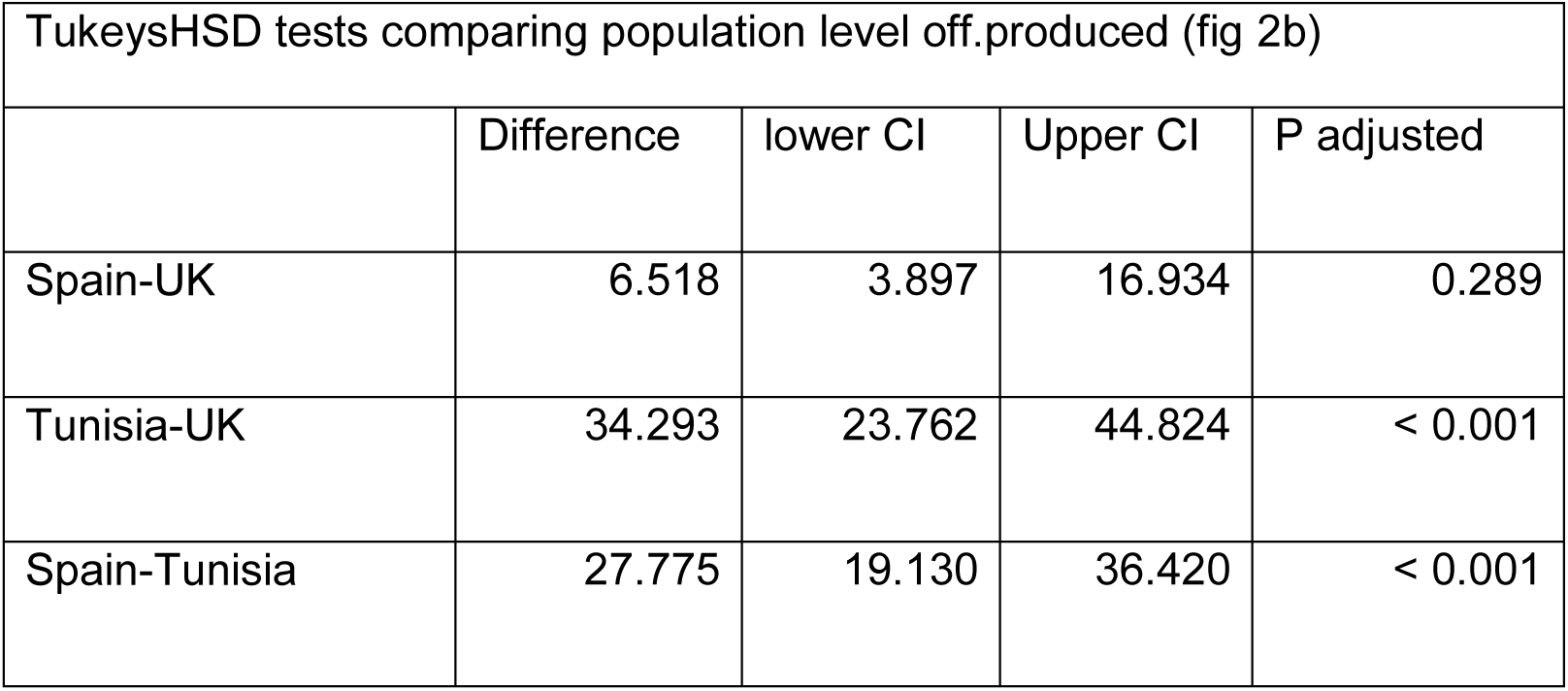
Supplementary table 3 showing differences in the offspring produced by SR*s* males when introgressed onto 39 isolines across three populations (Spain n=16, Tunisia n=15, UK n=8). Tukey’s post-hoc tests were used to test how the populations differ from each other. The mean number of offspring produced for each isofemale line was calculated using 20-40 replicates.

| TukeysHSD tests comparing population level off.produced (fig 2b) |  |  |  |  |
| --- | --- | --- | --- | --- |
|  | Difference | lower CI | Upper CI | P adjusted |
| Spain-UK | 6.518 | 3.897 | 16.934 | 0.289 |
| Tunisia-UK | 34.293 | 23.762 | 44.824 | < 0.001 |
| Spain-Tunisia | 27.775 | 19.130 | 36.420 | < 0.001 |

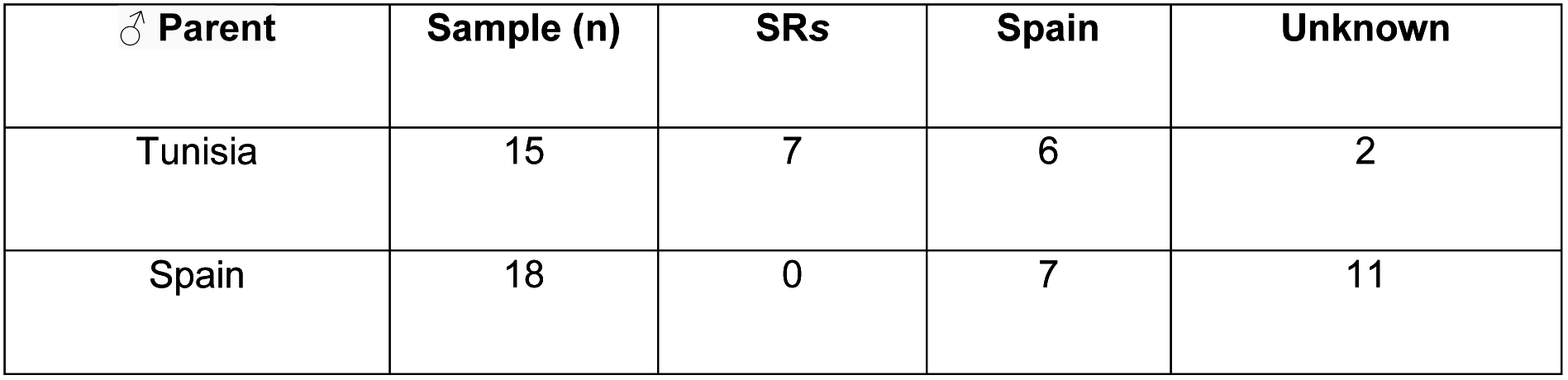
Supplementary table 4 shows the X-chromosome status of males produced by backcrossing hybrid females carrying one SR*s* and one Spanish X-chromosome to either a Tunisian male, to test for the rescue of the SR*s* phenotype, or to a Spanish male. The backcross to a Tunisian male confirms rescue of the SR*s* phenotype in 7 males. Male types were classified based upon the sex-ratio of the offspring produced as follows: SR*s* if the sex ratio of their offspring was >85% female, non-driving if the sex ratio was 50:50, and unknown if they produced 5 or fewer offspring.

Supplementary table 5 Differences between three populations in the proportion of offspring that are female when an SR*s* males was introgressed onto an isoline from that population. Number of isofemale lines used differed across three populations (Spain n=16, Tunisia n=15, UK n=8). Tukey’s post-hoc tests were used to test how the populations differ from each other. The mean proportion of female offspring was calculated for each isofemale line based on 20-40 replicate introgressed males from that isoline.

| TukeysHSD tests comparing population level suppression for (fig 5) |  |  |  |  |
| --- | --- | --- | --- | --- |
|  | Difference | lower CI | Upper CI | P adjusted |
| Spain-UK | 0.00682 | 0.042 | 0.0556 | 0.937 |
| Tunisia-UK | 0.08567 | 0.135 | 0.0363 | < 0.001 |
| Spain-Tunisia | 0.09250 | 0.133 | 0.0519 | < 0.001 |

## References

Babcock, C. S. & Anderson, W. W. 1996. Molecular evolution of the sex-ratio inversion complex in Drosophila pseudoobscura: Analysis of the esterase-5 gene region. Molecular Biology and Evolution, 13, 297–308.

Balanya, J., Oller, J. M., Huey, R. B., Gilchrist, G. W. & Serra, L. 2006. Global genetic change tracks global climate warming in Drosophila subobscura. Science, 313, 1773–1775.

Bastide, H., Cazemajor, M., Ogereau, D., Derome, N., Hospital, F. & Montchamp-Moreau, C. 2011. Rapid Rise and Fall of Selfish Sex-Ratio X Chromosomes in Drosophila simulans: Spatiotemporal Analysis of Phenotypic and Molecular Data. Molecular Biology and Evolution, 28, 2461–2470.

Burt, A. & Trivers, R. 2006. Genes in conflict: The biology of selfish genetic elements, Cambridge, Massachusetts, Harvard University Press.

Cobbs, G. 1992. Sex chromosome loss induced by the “sex-ratio” trait in Drosophila pseudoobscura males. J Heredity, 83, 81–84.

Crespi, B. & Nosil, P. 2013. Conflictual speciation: species formation via genomic conflict. Trends in Ecology & Evolution, 28, 48–57.

David, J., Gibert, P., Legout, H., PÉTavy, G., Capy, P. & Moreteau, B. 2005. Isofemale lines in Drosophila: an empirical approach to quantitative trait analysis in natural populations. Heredity, 94, 3–12.

Dyer, K. A. 2012. Local selection underlies the geographic distribution of sex-ratio drive in Drosophila neotestacea. Evolution, 66, 973–984.

Dyer, K. A., Charlesworth, B. & Jaenike, J. 2007. Chromosome-wide linkage disequilibrium as a consequence of meiotic drive. Proceedings of the National Academy of Sciences of the United States of America, 104, 1587–1592.

Hamilton, W. D. 1967. Extraordonary Sex-Ratios. Science, 156, 477–488.

Hauschteckjungen, E. 1990. Postmating reproductive isolation and modification of the sex-ratio trait in Drosophila subobscura induced by the sex-chromosome gene arrangement A^2+3+5+7^. Genetica, 83, 31–44.

Herrig, D., Modrick, A., Brud, E. & Llopart, A. 2014. Introgression in the Drosophila subobscura--D. madeirensis sister species: evidence of gene flow in nuclear genes despite mitochondrial differentiation. Evolution, 68, 705–719.

Holman, L., Freckleton, R. P. & Snook, R. R. 2008. What use is an infertile sperm? A comparative study of sperm-heteromorphic Drosophila. Evolution, 62, 374–85.

Jaenike, J. 2001. Sex chromosome meiotic drive. Annual Review of Ecology and Systematics, 32, 25–49.

Johnson, N. 2010. Hybrid incompatibility genes: remnants of a genomic battlefield? Trends Genet, 26, 317–325.

Jungen, H. 1967. Abnormal sex ratio, linked with inverted gene sequence, in populations of D. subobscura from Tunisia. Drosophila Information Service, 42, 109.

Jungen, H. 1968. Inversion polymorphism in Tunisian populations of Drosophila subobscura Collin. Archiv der Julius Klaus-Stiftung für Vererbungsforschung, Sozialanthropologie und Rassenhygiene, 43, 53–55.

Krimbas, C. B. 1993. Drosophila subobscura, Biology, genetics and inversion polymorphisms. Hamburg, Verlag Dr. Kovac.

Lenington, S., Franks, P. & Williams, J. 1988. Distribution of T-Haplotypes in Natural Populations of Wild House Mice. Journal of Mammalogy, 69, 489–499.

Lindholm, A. K., Dyer, K. A., Firman, R. C., Fishman, L., Forstmeier, W., et al. 2016. The Ecology and Evolutionary Dynamics of Meiotic Drive. Trends Ecol Evol, 31, 315–26.

Manser, A., Lindholm, A. K., Konig, B. & Bagheri, H. C. 2011. Polyandry and the decrease of a selfish genetic element in a wild house mouse population. Evolution, 65, 2435–2447.

Markow, T. A. & O’grady, P. 2005. Drosophila: a guide to species identification and use, Academic Press.

Mcdermott, S. R. & Noor, M. A. F. 2012. Mapping of within-species segregation distortion in Drosophila persimilis and hybrid sterility between D. persimilis and D. pseudoobscura. Journal of Evolutionary Biology, 25, 2023–2032.

Phadnis, N. & Orr, H. A. 2009. A Single Gene Causes Both Male Sterility and Segregation Distortion in Drosophila Hybrids. Science, 323, 376–379.

Pinzone, C. A. & Dyer, K. A. 2013. Association of polyandry and sex-ratio drive prevalence in natural populations of Drosophila neotestacea. Proceedings of the Royal Society B-Biological Sciences, 280, 8.

Prevosti, A. 1974. Chromosomal inversion polymorphism in the southwestern range of Drosophila subobscura distribution area. Genetica, 45, 111–124.

Prevosti, A., De Frutos, R., Alonso, G., Latorre, A., Monclus, M. & Martinez, M.-J. 1984. Genetic differentiation between natural populations of Drosophila subobscura in the Western Mediterranean Area with respect to chromosomal variation. Genet. Sel. Evol 16, 143–156.

Price, T. A. R., Bretman, A., Gradilla, A. C., Reger, J., Taylor, M. L., Giraldo-Perez, P., et al. 2014. Does polyandry control population sex ratio via regulation of a selfish gene? Proceedings of the Royal Society B-Biological Sciences, 281, 8.

Price, T. A. R., Hodgson, D. J., Lewis, Z., Hurst, G. D. D. & Wedell, N. 2008. Selfish Genetic Elements Promote Polyandry in a Fly. Science, 322, 1241–1243.

Sole, E. R. Balanya, J. Sperlich, D. & Serra, L. 2002. Long-term changes in the chromosomal inversion polymorphism of Drosophila subobscura. I. Mediterranean populations from Southwestern Europe. Evolution, 56, 830–835.

Stalker, H. D. 1961. Genetic systems modifying meiotic drive in Drosophila paramelanica. Genetics, 46, 177–202.

Sturtevant, A. H. & Dobzhansky, T. 1936. Geographical distribution and cytology of "sex ratio" in Drosophila pseudoobscura and related species. Genetics, 21, 473–490.

Sutter, A. & Lindholm, A. K. 2015. Detrimental effects of an autosomal selfish genetic element on sperm competitiveness in house mice. Proceedings of the Royal Society B-Biological Sciences, 282, 8.

Core Team, R, 2011. R: A Language and Environment for Statistical Computing, R Foundation for Statistical Computing, R Foundation for Statistical Computing.

Verspoor, R., Cuss, M. & Price, T. 2015. Age-based mate choice in the monandrous fruit fly Drosophila subobscura. Anim Behav, 102, 199–207.

Verspoor, R. L., Hurst, G. D. D. & Price, T. A. R. 2016. The ability to gain matings, not sperm competition, reduces the success of males carrying a selfish genetic element in a fly. Animal Behaviour, 115, 207–215.

